# Morphological responses of a temperate salt marsh foraminifer, *Haynesina* sp., to coastal acidification

**DOI:** 10.1101/2025.01.07.631753

**Authors:** Chris Powers, Alberto Paz, Amaelia Zyck, Kaylee Harri, Madison Geraci, Joan M. Bernhard, Ying Zhang

## Abstract

Coastal acidification leads to widespread impacts on calcifying organisms across the world’s oceans, which could result in decreased calcium carbonate deposition and the dissolution of calcium carbonate. As an abundant group of calcifying organisms, some protists within the phylum Foraminifera demonstrate potential success under elevated partial pressure of carbon dioxide (*p*CO_2_) due to their ability to modulate intracellular pH. However, little is known about their responses under more extreme acidification conditions that are already seen in certain coastal environments. Here we exposed *Haynesina*, a foraminiferal genus that is prevalent in temperate coastal salt marshes, to moderate (*p*CO_2_ = 2386.05+/-97.14 μatm) and high acidification (*p*CO_2_ = 4797.64+/-157.82 μatm) conditions through the duration of 28 days. We demonstrate that although this species is capable of withstanding moderate levels of coastal acidification with little impact on their overall test thickness, they could experience deposition deficiency and even dissolution of the calcareous test under highly elevated *p*CO_2_. Interestingly, such a deficit was primarily seen among live foraminifera, as compared to dead specimens, throughout the four-week experiment. We propose that a combination of environmental stress and the physiological process of test formation (i.e., calcite precipitation) could induce thinning of the test surface. Therefore, with the acceleration of coastal acidification due to anthropogenic production of CO_2_, benthic foraminifera amongst coastal ecosystems could reach a tipping point that leads to thinning and dissolution of their calcareous tests, which in turn, will impair their ecological function as a carbon sink.

**Importance:** The calcareous foraminifera protists are responsible for large proportions of calcium carbonate production across the global ocean. Their responses to ocean and coastal acidification are essential for understanding carbon and mineral cycling in diverse marine ecosystems. However, relatively few studies have examined more extreme conditions related to what is seen in coastal habitats (e.g. *p*CO_2_ > 2,500 μatm), and the response of individual test chambers have never been inspected. Here, we consider the response of *Haynesina* sp., a benthic foraminifera obtained from temperate coastal sediments to moderate and high acidification regimes. Comparison of test thicknesses across treatment conditions and among individual chambers of *Haynesina* sp. revealed potential tolerance under moderate acidification but demonstrated impaired new chamber formation and test dissolution under high acidification. Our results suggest that with growing anthropogenic CO_2_ production, foraminifera could reach a tipping point that leaves their ecological function as a carbon sink at greater risk.

## Introduction

Increasing anthropogenic production of CO_2_ into the atmosphere has resulted in rising ocean temperatures, causing increases in sea levels and changes in the ocean’s chemistry (1). As excess CO_2_ dissolves in seawater, the concentration of hydrogen ions increases, leading to acidification and reduced carbonate ion concentration (2–4). Anthropogenic-driven acidification has resulted in decreasing pH across the world’s oceans, with drops of 0.1 to 0.3 possible in open ocean waters within the next 100 years (1). These changes are exacerbated in coastal areas, such as Narragansett Bay (RI, USA) and Long Island Sound (NY-CT-RI, USA), where low pH (< 7.2), high *p*CO_2_ (> 2,500 μatm), and aragonite undersaturation (Ω_aragonite_ < 1) have been observed periodically in bottom waters (5).

Increased *p*CO_2_ may have dire impacts for organisms in the ocean, such as those that produce their own shells, tests, or skeletons by depositing calcium carbonate minerals (*i.e.*, calcite and aragonite). As *p*CO_2_ of seawater increases, the aragonite saturation (Ω_aragonite_) and calcite saturation (Ω_calcite_) state of seawater decreases, undermining the shell formation of marine organisms (1, 6, 7). However, the impact of ocean acidification on individual calcifying taxa remains difficult to generalize due to the confounding effects of elevated *p*CO_2_ and organisms’ own acclimatory responses (2, 8). Such phenomena call for more studies on specific taxa to gain a more complete picture of their responses to acidification.

The Foraminifera is a phylum of unicellular microeukaryotes prevalent in marine and sedimentary ecosystems, from shallow to deep water depths. Many foraminifera produce calcareous tests through calcite precipitation (6, 9). Foraminifera account for an estimated 25% of calcium carbonate deposition across the world’s oceans, attesting to their significant roles in carbon and mineral cycling (10). The test morphology within certain foraminifera genera varies depending on their reproductive stage (11–17). The sexually reproducing microspheric foraminifera are characterized by a relatively small proloculus (i.e., the first chamber formed during growth) due to cytoplasmic requirements of progeny produced through gametogenesis and fertilization (11). In contrast, the asexually reproducing megalospheric foraminifera have a relatively large proloculus and inherit significant volumes of the parental cytoplasm, including potential symbionts (11). Additionally, the microspheric foraminifera tend to form significantly more chambers than their megalospheric counterparts while exhibiting heteromorphic test structure (12, 13).

Despite their abundance in temperate coastal systems, most acidification studies of foraminifera focus on species from reef-associated systems or the open ocean (**Supplemental Table S1**). A few studies examine the responses of individual foraminifera to coastal acidification, but more extreme conditions related to what is seen in coastal habitats (e.g. *p*CO_2_ > 2,500 μatm) are rarely considered. Common strategies for measuring foraminifera responses to acidification involve tracking chamber formation rates and surface morphological changes, with prior studies demonstrating variable influences of acidification on different species of foraminifera (18–31). The direct measurement of test thickness, however, has not been systematically applied to study the acidification responses in foraminifera.

In this study, laboratory treatments were conducted to examine the response of *Haynesina* sp., a benthic foraminifer identified from coastal sediments in Rhode Island, USA, to moderate (*p*CO_2_ = 2386.05+/-97.14 μatm) and high acidification (*p*CO_2_ = 4797.64+/-157.82 μatm) conditions. Thicknesses of foraminifera tests were mapped using X-ray tomography, enabling systematic comparisons throughout individual test chambers. The treatments were applied to both live and dead foraminifera, which provides an opportunity for examining the biotic and abiotic responses of foraminifera and their calcareous remains to coastal acidification.

## Results

### Acidification challenges

Specimens were picked from surface sediments obtained from a mudflat associated with the Quonochontaug Salt Marsh (41.336824, -71.72107) on June 19, 2023 and September 26, 2023, respectively, for two replicate pH manipulation experiments (**Table 1**). Each replicate trial was performed over a period of 28 days with three treatment tanks (**Figure 1A**). The target pH of each tank was maintained at 8.1, 7.6, and 7.2, respectively, representing the open ocean pH (no *p*CO_2_ manipulation), a middling condition (moderately elevated *p*CO_2_), and the lowest pH (highly elevated *p*CO_2_) that simulates stress events previously observed in Long Island Sound and Narragansett Bay (Wallace et al. 2014).

**Figure 1.**
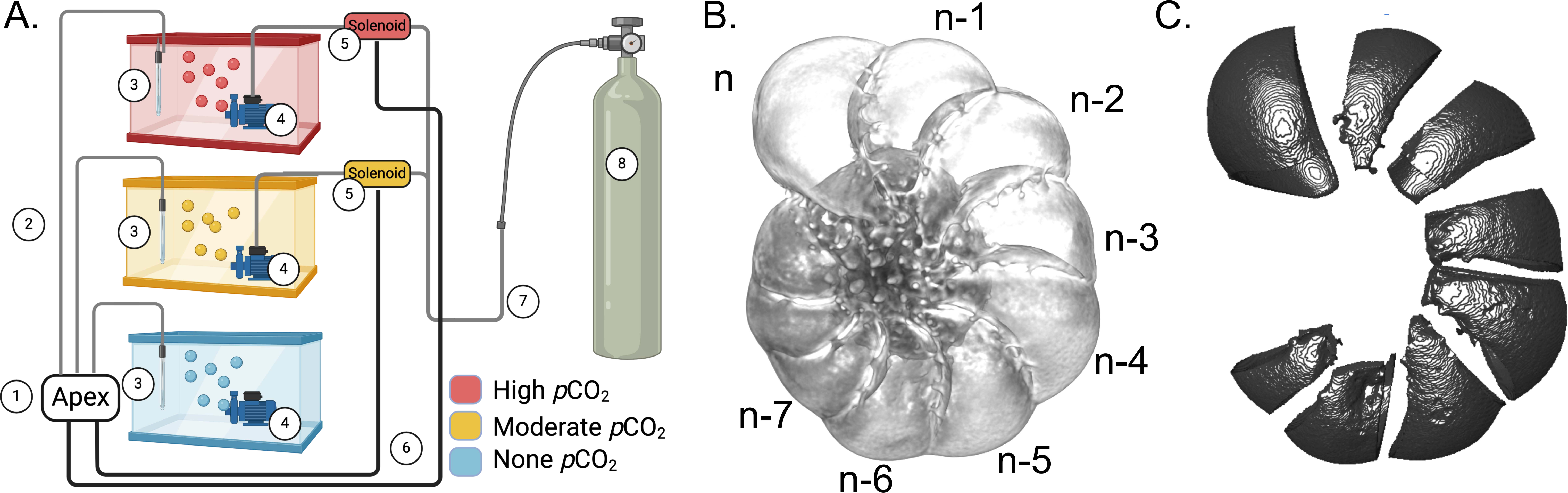
(A) Schematic representation of the experimental setup for *p*CO_2_ manipulation. Components of the diagram are as follows: ① the Apex controller system, ② wires connecting pH probe to APEX controller, ③ pH probe, ④ water pumps with Venturi injector, ⑤ solenoids controlling gas flow for the elevated *p*CO_2_ treatments, ⑥ wires connecting the apex controller to the solenoids, ⑦ gas tubing connecting the CO_2_ gas supply to the treatment tanks, ⑧ CO_2_ gas supply. (B) A foraminifera test with labels showing the 8 newest chambers. (C) An example of isolated exteriorly facing test areas from each chamber for test-thickness analysis.

**Table 1.** Carbonate system parameters measured from laboratory acidification experiments. Values shown are averaged across all timepoints taken throughout the four-week period. Variation is shown in terms of the standard error of the mean. pH_T_ represents the pH on the total scale, *p*CO_2_ is the partial pressure of carbon dioxide, TA is the total alkalinity, and Ω_calcite_ is calcite saturation. All represents the combined data for Summer 2023 and Fall 2023.

| Trial | Parameter | Tank 1 | Tank 2 | Tank 3 |
| --- | --- | --- | --- | --- |
| All | $\text{pH}_T$ | 7.101±0.016 | 7.376±0.015 | 7.826±0.017 |
| | $p\text{CO}_2$ ($\mu\text{atm}$) | 4797.64±157.82 | 2386.05±97.14 | 431.18±18.90 |
| | TA ($\mu\text{mol/kg}$) | 2511.68±55.32 | 2400.87±44.86 | 1395.91±39.82 |
| | $\Omega_{\text{calcite}}$ | 0.843±0.041 | 1.498±0.057 | 2.153±0.121 |
| Summer 2023 | $\text{pH}_T$ | 7.119±0.026 | 7.326±0.016 | 7.884±0.016 |
| | $p\text{CO}_2$ ($\mu\text{atm}$) | 4495.21±222.72 | 2752.34±104.95 | 400.31±19.36 |
| | TA ($\mu\text{mol/kg}$) | 2459.07±83.36 | 2486.00±70.98 | 1522.837 |
| | $\Omega_{\text{calcite}}$ | 0.865±0.069 | 1.380±0.075 | 2.594±0.138 |
| Fall 2023 | $\text{pH}_T$ | 7.084±0.016 | 7.427±0.016 | 7.764±0.018 |
| | $p\text{CO}_2$ ($\mu\text{atm}$) | 5100.08±197.32 | 2019.77±77.09 | 464.61±31.38 |
| | TA ($\mu\text{mol/kg}$) | 2564.289±<br>73.071 | 2315.745±<br>46.270 | 1258.398±<br>32.937 |
| | $\Omega_{\text{calcite}}$ | 0.821±0.045 | 1.615±0.075 | 1.674±0.060 |

Seawater chemistry of the treatment tanks was measured tri-weekly to monitor total scaled pH (pH_T_), *p*CO_2_ (μatm), total alkalinity (TA), and calcite saturation state (Ω_calcite_) (**Supplemental Table S2**). The experimental treatments resulted in three distinct *p*CO_2_ regimes: no *p*CO_2_ manipulation (431.18+/-18.90 μatm), moderately elevated *p*CO_2_ (2386.05+/-97.14 μatm), and highly elevated *p*CO_2_ (4797.64+/-157.82 μatm). Additionally, untreated control samples were collected following field sampling and before laboratory treatment (**Materials and Methods**). Measurements of the calcite saturation state indicated supersaturation (Ω_calcite_ > 1) under both non-elevated (Tank 3) and moderately elevated (Tank 2) *p*CO_2_ treatments. However, calcite undersaturation (Ω_calcite_ < 1) was observed under the highly elevated (Tank 1) *p*CO_2_ treatment (**Table 1**).

### Microscopy and three-dimensional test reconstruction

Three-dimensional (3D) reconstruction of foraminifera tests was achieved with a voxel size of 0.57 µm (resolution around 1 µm) using microCT scanning (**Figure 1B**). Individual chambers were extracted during image processing following the 3D reconstruction, and the thicknesses of exteriorly facing test areas were measured (**Figure 1C**, **Materials and Methods**). From each replicate trial, we initially collected at least 6 live and 6 dead specimens from each treatment tank, and at least 8 specimens as untreated controls. Some tests were lost during handling, and some others were damaged when mounted for microCT scanning due to their delicate nature. These structurally damaged tests were found among all treatment conditions and were not included in further analyses (**Supplemental Table S3**).

### Assignment of test morphology

Foraminifera specimens collected from each treatment were classified into two distinct groups, microspheric and megalospheric, based on their heteromorphic test geometry (**Materials and Methods**). A total of 18 megalospheric and 58 microspheric specimens were identified across all treatments based on a bimodal distribution of proloculus sizes (**Supplemental Figure S1**). These assignments were independently verified by examining the number of chambers for all assigned tests (**Figure 2A**), as the proloculus size and number of chambers are both known to vary greatly between sexual and asexual reproductive stages (13, 16).

**Figure 2.**
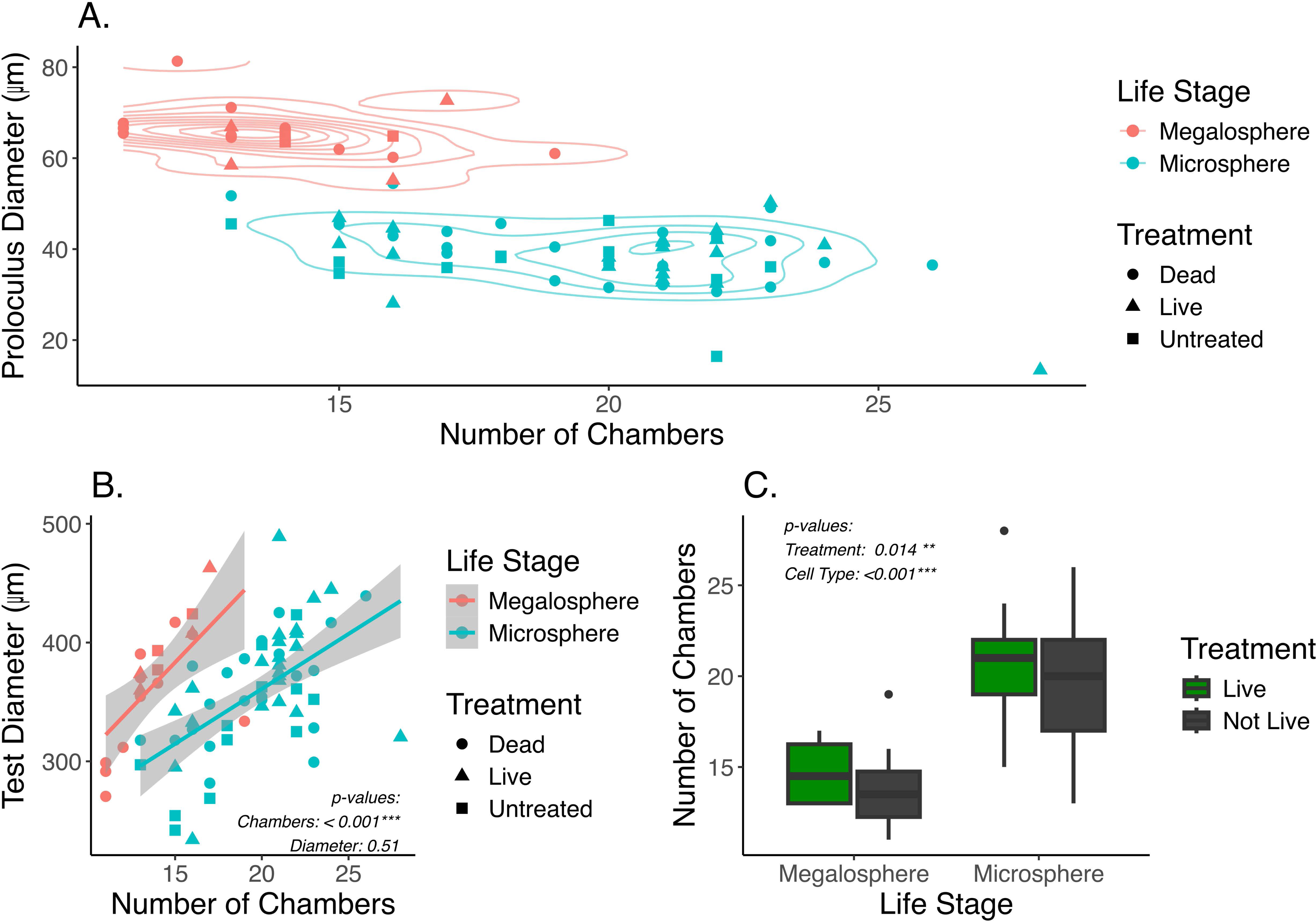
(A) Plot showing the number of chambers against the proloculus diameter. Each point represents a foraminifer. Color represents the assignment of two life stages, microsphere or megalosphere, based on the diameter of proloculus. Symbol shape represents different treatment groups. (B) Comparison of the number of chambers and the test diameter between two life stages. P-values are based on one-way ANOVA accounting for different life stages (Materials and Methods). (C) Box and whisker plot showing distribution of the number of chambers in live and not-live foraminifera between the two life stages. The “not-live” specimens include both untreated and dead treated samples. P-values are based on two-way ANOVA accounting for life stages and live vs. not-live treatment groups (Materials and Methods).

The mean proloculus diameter of the megalospheric tests (65.47+/-5.77 µm) was approximately 70% larger than that of the microspheric tests (38.78+/-7.12 µm), consistent with morphological features known for these two life stages (**Supplemental Table S4**). The number of chambers per microspheric test was significantly higher than per megalospheric test (One-Way ANOVA: F_1,74_=50.28, p < 0.001). However, the overall test diameter was comparable between the microspheric and megalospheric specimens (**Figure 2B**). A significant increase in the number of chambers was seen when live versus not-live (including untreated and dead treated) specimens were compared (Two-Way ANOVA: F_1,72_=6.319, p = 0.0014) (**Figure 2C**). This likely resulted from growth of live foraminifera through the duration of the four-week incubation period. For microspheric foraminifera, the average number of chambers in the live and not-live groups are 20 and 19, respectively. For megalospheric foraminifera, the average number of chambers in the live and not-live groups are 15 and 14, respectively. Therefore, a putative growth of about one chamber was expected among the live foraminifera compared to their untreated or dead counterpart (**Supplemental Table S4**). However, the TukeyHSD comparison of live and not-live foraminifera did not appear to support the statistical significance within each of the two different life stages. This indicates a high level of variability in the number of chambers among individual foraminifera.

### Variation of test thicknesses across different treatments

The test morphology was not assigned until after the experimental period due to challenges in keeping foraminifera alive following microCT scanning, which involves bleaching to remove soft tissues and the exposure to high X-rays through extended scanning period during imaging (**Materials and Methods**). Therefore, the experimental treatments had an uneven number of megalospheric and microspheric specimens. Due to the sparsity of megalospheric samples in multiple treatment conditions (**Supplemental Table S3**), statistical comparison of test thicknesses across different treatment groups (e.g. treated versus untreated, different *p*CO_2_ treatment conditions, or live versus dead treatments) were performed only with the microspheric specimens.

Comparisons of test thicknesses indicated substantial variations among individual foraminifera. The effect size related to the individual variance in two-way ANOVA analyses was around 0.283-0.365, which is 1-2 orders of magnitude higher than what was seen in the effect of experimental treatments (**Table 2**). Although a relatively small effect was seen in the factor that compared different treatments to the no-treatment control, they revealed variable responses. The non-elevated and moderately elevated *p*CO_2_ treatments had negligible effect sizes (η^2^ ≤ 0.01) compared to the untreated control. In contrast, specimens in the highly elevated *p*CO_2_ treatments had thinner tests compared to the untreated control, with effect sizes of 0.023 and 0.014, respectively, for the live and dead foraminifera (**Table 2**).

**Table 2.** Comparison of test thicknesses between each treatment to the untreated control. Effect size (η^2^) was calculated using two-way ANOVA models that account for both the treatment factor and the variations among individual foraminifera.

| Treatments ($p\text{CO}_2$) | Live treatment | | Dead treatment | |
| --- | --- | --- | --- | --- |
|  | Treated vs. Untreated | Individual variance | Treated vs. Untreated | Individual variance |
| Non-Elevated | 0.001 | 0.283 | 0.002 | 0.312 |
| Moderately Elevated | 0.005 | 0.300 | 0.000 | 0.313 |
| Highly Elevated | 0.023 | 0.365 | 0.014 | 0.293 |

Significant differences in test thicknesses were observed for both live and dead specimens across the different *p*CO_2_ treatments, where a slightly higher effect size was observed among the live (η^2^ = 0.024) than the dead treatments (η^2^ = 0.011) (**Figure 3A-B**). Specifically, thinner tests were observed in the live cell treatments under highly and moderately elevated *p*CO_2_ compared to the non-elevated *p*CO_2_ (**Figure 3A**). Differential responses between live and dead treatments were also observed in the distribution of test thicknesses. The largest effect was seen in the highly elevated *p*CO_2_ treatment (η^2^ = 0.072), showing significant thinning of tests in live compared to dead treated foraminifera (**Figure 3C**), while the non-elevated and moderately elevated *p*CO_2_ treatments had little evidence of thinning when comparing the live and dead treatments (**Figure 3D-E**). Comparisons on each of the eight newest chambers also revealed significantly thinner tests among the live compared to the dead foraminifera, particularly, under the high *p*CO_2_ treatment (**Figure 4**). Interestingly, higher effect sizes (η^2^ > 0.1) were observed in the six newest chambers (from n to n-5), while a lower effect (η^2^ < 0.1) was seen in chambers n-6 and n-7 among the high *p*CO_2_ treatment of live versus dead specimens (**Figure 4A**).

**Figure 3.**
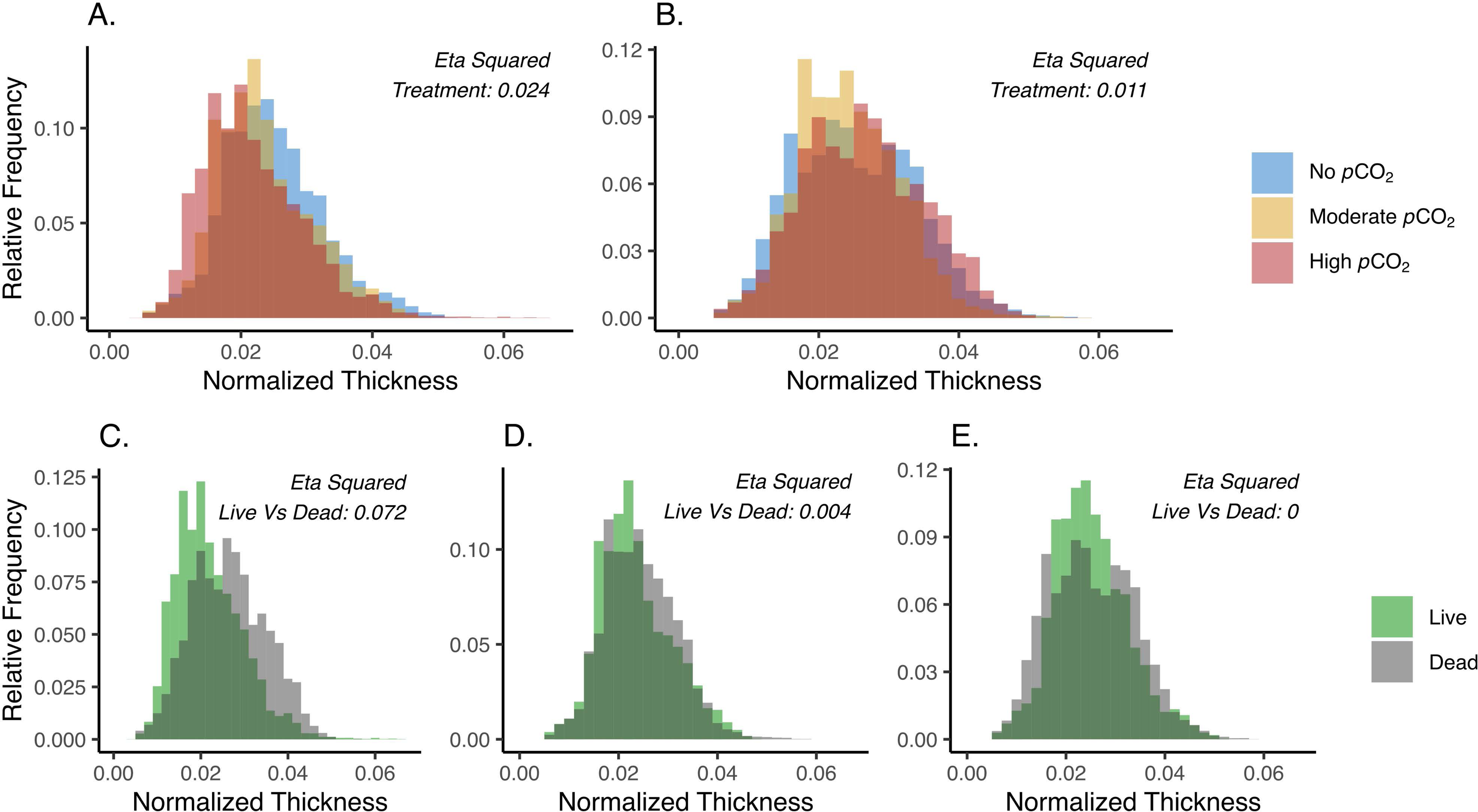
(A-B) Distribution of normalized test thickness across the highly-(red), moderately-(gold), and no-(Blue) elevated *p*CO_2_ conditions among the live (A) and dead (B) specimens. (C-E) Distribution of the normalized test thickness between the live (green) and dead (gray) foraminifera at the highly elevated pCO2 (C), moderately elevated pCO2 (D), and no elevated pCO2 (E) treatments. Only Microspheric foraminifera were used in this analysis (Materials and Methods). Untreated specimens were not included in this comparison. The η^2^ values are effect sizes derived from two-way ANOVA (Materials and Methods).

**Figure 4.**
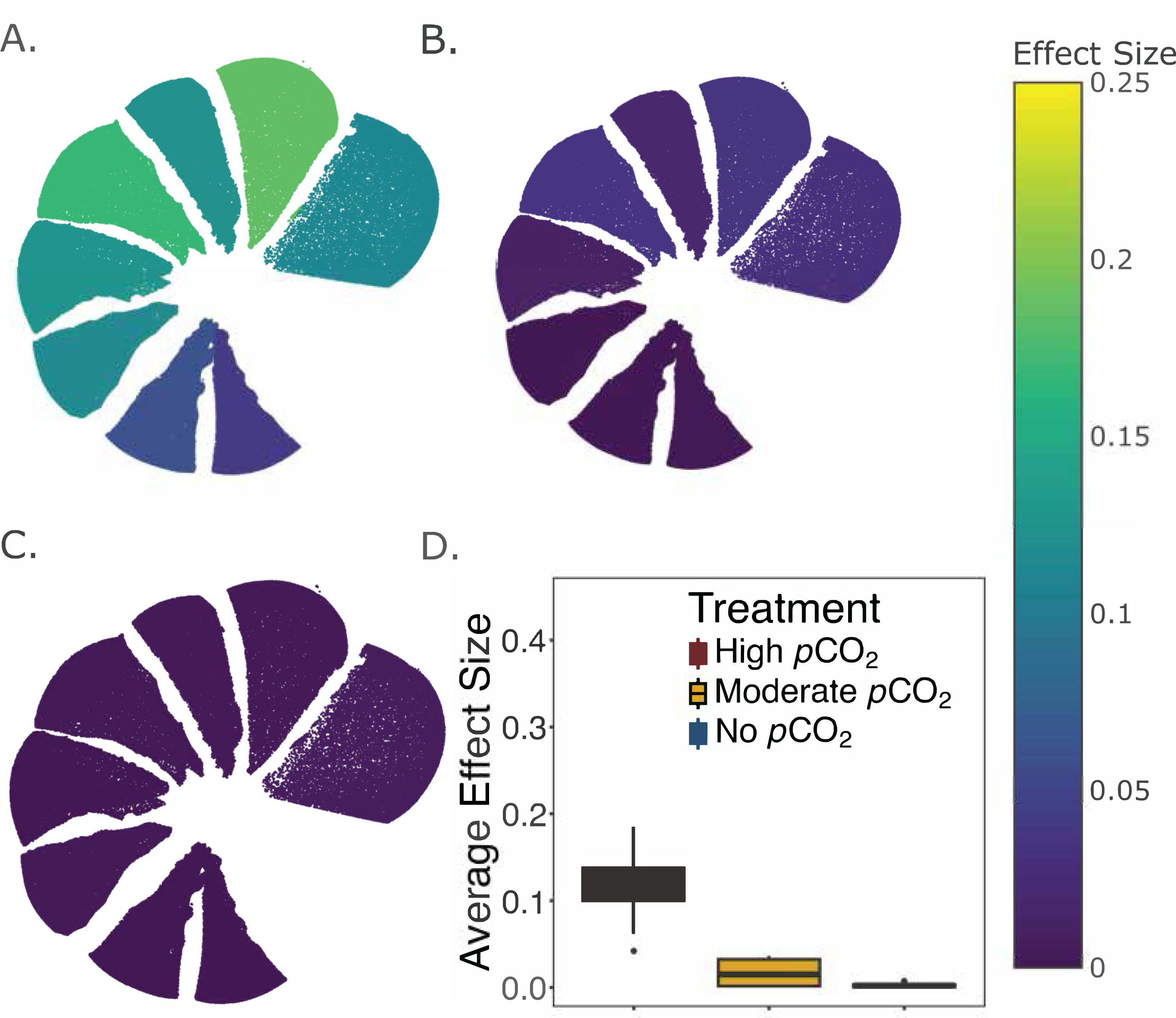
(A-C) Comparison of test thicknesses between live and dead treatments within each of the 8 newest chambers under the highly elevated pCO2 (A), moderately elevated pCO2 (B), and no elevated pCO2 (C) treatment conditions. Untreated specimens were not included in this comparison. The color of each chamber represents the effect size (η^2^) of the live vs. dead factor in a two-way ANOVA that accounts for variations among individual foraminifera. (D) Box and whisker plot showing the median effect size and the first and third quartiles across all 8 chambers for each treatment. Only microspheric foraminifera were used in this analysis (Materials and Methods). The total number of specimens in each treatment group is documented in Supplemental Table S3.

## Discussion

Calcareous foraminifera serve as carbon sinks across the global ocean by incorporating calcium carbonate to their tests, sequestering carbon from the surrounding seawater. Benthic foraminifera play significant roles in the worldwide carbon budget with an estimated production of 200 million tons of calcium carbonate per year (10). However, calcium carbonate production by foraminifera could be negatively impacted by the anthropogenic production of excess CO_2_, which causes ocean and coastal acidification, subsequently decreasing the saturation of carbonate system in the marine environment.

Ocean and coastal acidification could have mixed impacts on foraminifera, with studies noting that some foraminifera species can survive in moderate elevation of *p*CO_2_ (790 - 1865 μatm) without major growth defects (19, 20) or even showing increased growth rates (24). However, the majority of studies indicate either decreased growth rate or defects in morphology at decreased pH (7.4 - 7.9) or at moderate to highly elevated *p*CO_2_ (e.g. up to 3247 μatm) (18–30). Study of *Haynesina germanica,* a temperate salt marsh foraminifer closely related to the *Haynesina* sp. examined in this study, suggests their feeding-related test ornamentation can be deformed during prolonged treatments (36 weeks) of moderately elevated *p*CO_2_ (380-1000 ppm) (30). However, morphological alteration has not been systematically documented throughout the entire test. Further, with the projected increases of ocean *p*CO_2_, more extreme acidification conditions, such as those observed in porewaters of estuarine mudflat sediments (32), will become more impactful to coastal foraminifera.

To our knowledge, this is the first study that differentiates the two alternative generations of the foraminifera lifecycle, microsphere and megalosphere, in examining foraminifera responses to acidification. This distinction could be crucial as varied test structures have been observed between the two life stages of foraminifera, such as those documented in some Elphidiids (13). This variability is shown in our experimental data, where microspheric and megalospheric foraminifera had varied test thickness distributions and different levels of sensitivity to laboratory treatments (**Supplemental Figure S2**). The classification of microspheric and megalospheric foraminifera was based on a bimodal distribution of proloculus diameters (**Supplemental Figure S1**). This assignment was independently verified by examining the number of chambers between these two life stages, where the microspheric foraminifera had a significantly higher number of chambers compared to the megalospheric foraminifera (**Figure 2**). Our current technology, however, supports the identification of life stages only after experimental treatments because of the destructive nature of extended exposure to high X-rays during MicroCT scanning (**Materials and Methods**). As a result, an insufficient number (n < 3) of megalospheres was included in some treatment conditions (**Supplemental Table S3**), and the test-thickness analyses were performed only on microspheres. Therefore, the response megalospheres to coastal acidification remains unknown, which could be a topic for future investigations.

Compared to the untreated group, the experimental treatment of both live and dead microspheric foraminifera had a larger effect size (η^2^ > 0.01) in highly elevated *p*CO_2_ relative to the little to no effect (η^2^ ≤ 0.005) in non- or moderately elevated *p*CO_2_ (**Table 2**). Most calcareous foraminifera form tests that are mainly composed of calcite, which is structurally more stable (33) and less prone to dissolution (34) than the calcium carbonate polymorph aragonite. Given that calcite oversaturation was measured in the moderate treatment (Ω_calcite_ = 1.498+/-0.057), it is unsurprising that *Haynesina* test thickness exhibited little to no change in the moderately elevated *p*CO_2_. In contrast, the highly elevated *p*CO_2_ treatment exhibited calcite undersaturation (Ω_calcite_ = 0.843+/-0.041), consistent with the observation of test thinning in both live and dead specimens (**Table 1 & Table 2**). It is worth noting that the treatment period of our study was 4 weeks, significantly shorter than the long-term treatment (36 weeks) performed on *Haynesina germanica* (30). Future studies are required to examine acidification responses through extended periods under both moderately and highly elevated *p*CO_2_, especially as such prolonged exposure becomes relevant to the coastal benthic environment.

Typically, new chambers in foraminifera precipitate via multiple steps: (1) formation of an outer organic layer, which is a protective envelope that defines the bound of the new chamber; (2) construction of the primary organic sheet, which forms under the protective envelope; and (3) calcification around the organic sheet (33, 35–38). The calcification relies on the maintenance of a local environment within the protective envelope with conditions favorable for calcium carbonate precipitation (36, 37, 39, 40). This process could be facilitated by vacuolar ATPases, which transport protons from the calcification site to vesicles that are then exported to the extracellular space (41). Therefore, maintenance of calcification-promoting conditions in foraminifera could involve potential energetic expenses due to consumption of ATP for proton export.

Comparing the acidification treatment of live versus dead specimens demonstrated significant differences in their responses to acidification, with the highest effect size observed in the live populations incubated under the high *p*CO_2_ condition (**Figure 3**). This indicates that thinning of foraminifera tests could be driven not only by calcite undersaturation, but also by the physiological activity of live foraminifera, likely related to the formation of new chambers (**Figure 2C**). The chamber-specific comparison of live and dead specimens has further emphasized the significant effect of foraminifera physiology on test chamber thickness under highly elevated *p*CO_2_. In particular, a more substantial effect size (η^2^ from 0.11 to 0.19) was observed in each of the six newest chambers (n to n-5) compared to chambers n-6 (η^2^ = 0.06) or n-7 (η^2^ = 0.04) (**Figure 4**), suggesting potential effects of new chamber formation in exacerbating test thinning in high *p*CO_2_ systems, likely due to the export of protons mediated by vacuolar ATPases.

Proton release during the formation of new test chamber can lead to increased proton concentration (**Figure 5A**), subsequently lowering the pH in the microenvironment that surrounds the foraminifera test (36). The decreasing pH alters calcite saturation (Ω_calcite_), which in turn can lead to potential dissolution of the test surface (**Figure 5B**). Under the no-elevated and moderately elevated *p*CO_2_ treatments performed in this study, Ω_calcite_ is relatively high, and hence the decrease of pH caused by calcification could have less effect. However, Ω_calcite_ in the highly elevated *p*CO_2_ was close to the value of 1 (**Figure 5C**), below which dissolution is expected due to calcite undersaturation. Therefore, even a slight decrease of pH could have significant effects on the foraminiferal test, not only increasing the energy demands in promoting calcification and new chamber formation, but also resulting in the dissolution of existing test surfaces.

**Figure 5.**
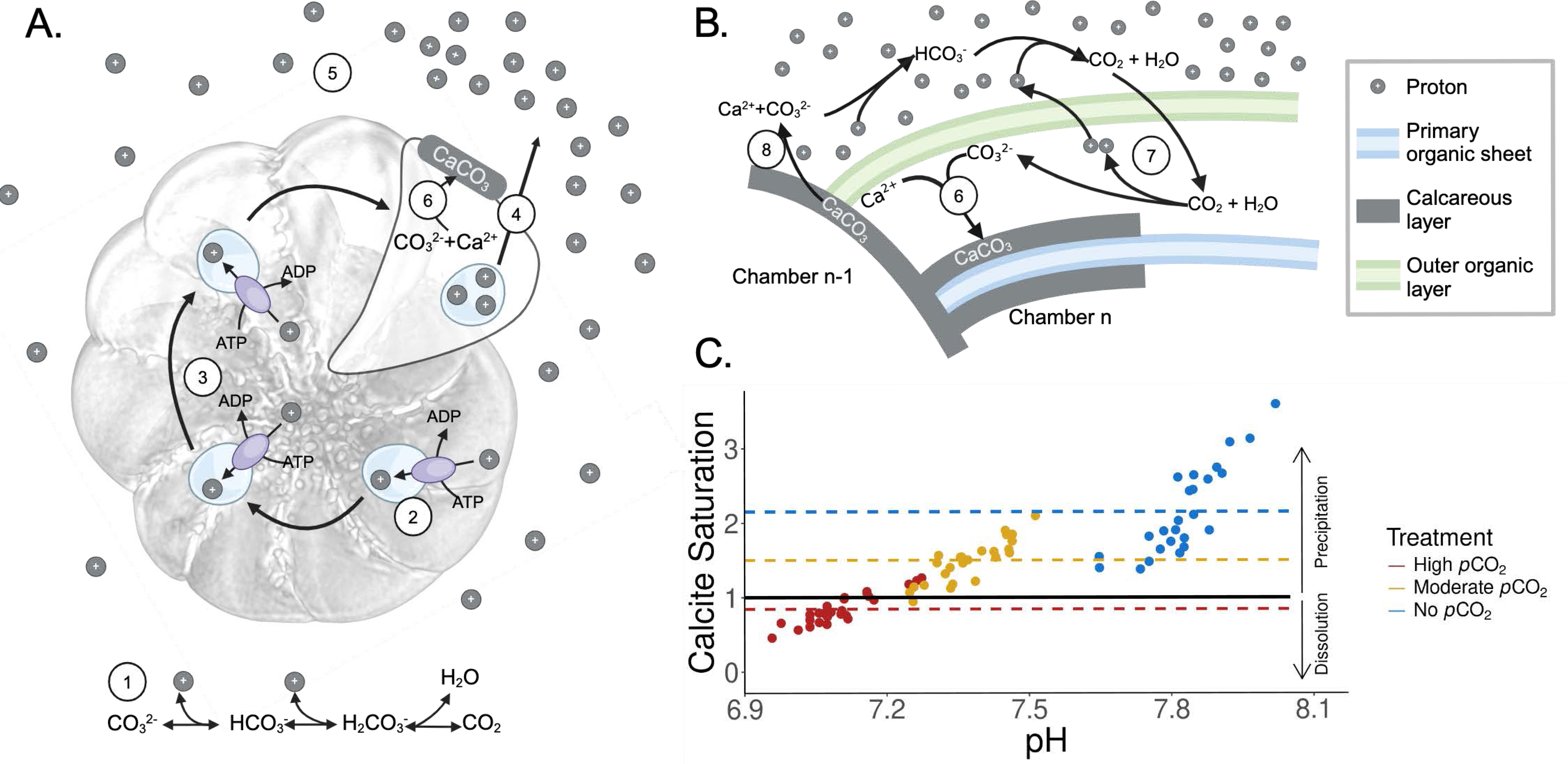
(A) Schematic of chamber formation in foraminifera. Components are as follows: ① the system of reactions dictating that increased CO_2_ results in increased proton concentration, ② vacuolar ATPases facilitate the export of protons by collecting them into vacuoles, as reported by (41), ③ proton vacuoles are moved throughout the cytoplasm to coordinate exocytosis, ④ protons are released through exocytosis, ⑤ protons diffuse outward and around the test, lowering pH in the microenvironment surrounding the actively growing foraminifera cell (36), ⑥ The proton-depleted environment allows for calcium carbonate precipitation. (B) Carbon chemistry during foraminiferal test formation. ⑦ foraminifera promote calcification through proton export, ⑧ test surface dissolution driven by acidification. (C) Calcite saturation state predicted based on tri-weekly experimental measurements acquired from this study. Each dot represents a measurement data point. Red represents the highly elevated *p*CO_2_ treatment, gold represents the moderately elevated *p*CO_2_ treatment, and blue represents the no elevated *p*CO_2_ treatment. The black horizontal line represents a calcite saturation of 1. Dashed lines represent the mean calcite saturation values of each treatment. Arrows on the right indicate the effect of calcite saturation state on the dissolution or precipitation of calcareous tests.

We suggest that newer chambers could be more sensitive to acidification than the older chambers, as the physiologically driven pH reduction is likely initiated in the extracellular space near the site of calcium carbonate precipitation of the new chamber (**Figure 5**). Our experimental observations of the *Haynesina* sp. (**Figure 4A**) support models of foraminifera calcification previously described in other studies (36, 41) and are consistent with observations from another foraminifera, *Ammonia* sp., where the lowest extracellular pH in its surrounding microenvironment was measured near the newest chamber (36, 42).

The ability of foraminifera to use proton pumping to manipulate carbonate chemistry is a competitive advantage against ocean and coastal acidification, as it enables the organism to decouple calcium carbonate precipitation from the chemistry of the surrounding seawater (36, 37, 40–42). However, our results suggest that in conditions near the borderline of calcite undersaturation, foraminifera could reach a tipping point that exacerbates the risk of test dissolution. Further, the energetic cost of proton pumping could increase with any continued rise of *p*CO_2_ (37), as foraminifera must overcome stronger concentration gradients to achieve an optimal calcification rate (43). This is notable, as the *p*CO_2_ conditions tested in this study have already been observed in coastal systems (5). Therefore, coastal benthic foraminifera are likely experiencing acidification stress that impairs new chamber formation and dissolves already formed test surfaces. With continued anthropogenic production of CO_2_, coastal acidification will accelerate in intensity and duration (44), leaving the ecological function of foraminifera as a carbon sink at greater risk.

## Materials and Methods

### Field sampling and sample preparation

Surface sediments were collected into 125-mL high density polyethylene Nalgene containers using a plastic scoop. Collected samples were sieved with USA standard sieves 120 (Thermo Fisher Scientific 039988.ON) and 40 (Thermo Fisher Scientific 039984.ON) to select for the size fraction between 125 µm and 425 µm. Isolated sediments were subsequently picked for approximately 600 specimens of *Haynesina* sp. using 50-µL calibrated pipettes (Drummond Scientific Company 2-000-050), which were pulled to a thin point over a bunsen burner, to isolate individual foraminifera while visualizing with a trinocular stereo microscope under 10-25x with maximum brightness (VanGuard 1372ZL). A subsample of 60 individuals were placed in 2 mL 6% sodium hypochlorite solution (Fisher Scientific NC1796686) for 12 hours to remove organic material from the test through bleaching. After bleaching, specimens were rinsed twice for five minutes with Milli-Q H_2_O (Type I H_2_O purified with EMD Millipore MilliQ EQ-7008). Eight of the bleached specimens were collected as a no-treatment control (i.e., untreated) and were retained in 100% ethanol at 4°C until microscopic imaging. The rest of the bleached specimens were used in the dead treatment and were stored in Milli-Q H_2_O at 4°C until experimental manipulation. The rest of the picked foraminifera were kept alive under room temperature in artificial seawater composed of Milli-Q water and Reef Pro Mix (Fritz Aquatics 80243) made at a salinity of 35 ppt until being used as live treatment in experimental manipulation.

### Experimental pCO_2_ manipulation

Experimental *p*CO_2_ manipulation was performed in three 75-liter glass treatment tanks with target pH maintained at 7.2 (Tank 1), 7.6 (Tanks 2), and 8.1 (Tank 3), respectively. All treatment aquaria were maintained with artificial seawater. Replicate foraminifera samples were introduced to treatment tanks in six-well plates sealed with 60-µm nylon mesh (Amazon ASIN#B092D8TJDQ). Each tank had 4 replicate six-well plates, with each plate contained 35 live foraminifera in one well and 3 bleached (dead) foraminifera in a separate well. The acidification treatments were designed following prior examples (45), with *p*CO_2_ levels controlled using an A3 Apex Aquarium Controller System (Bulk Reef Supply, SKU 251246). The Apex system measures pH and temperature (°C) every 10 seconds and adjusts the pH to a target value by injecting CO_2_ gas using controls of solenoid valves (**Figure 1A**). Three times per week (Monday, Wednesday, and Friday), 200 mL of tank water from each glass tank were filtered through 0.2 µm surfactant-free cellulose acetate (SFCA) syringe filters (Thermo Scientific 723-2520). This filtered seawater was stored at -20°C for stability before being used for carbonate system analysis (46). At each time point, pH was measured using a calibrated pH meter (OHAUS Aquasearcher 30589830), salinity was measured using a refractometer (Amazon ASIN#B018LRO1SU), and temperature was measured using the Apex controller temperature probe (Bulk Reef Supply, SKU 207517). Each Friday, the OHAUS pH meter was calibrated through examination of temperature and voltage correlation, and replicate wells of live foraminifera specimens were fed with *Skeletonema dohrnii* PA 250716_D1. The *S. dohrnii* was cultured in F/2 medium (47) under 12-hr light and 12-hr dark cycles. At the time of feeding, concentration of live *S. dohrnii* culture was quantified with a hemocytometer (Fisher Scientific 02-671-6) to determine the volume used for feeding live foraminifera. An average of 124 µL *S. dohrnii* culture was used in each feeding to add approximately 25,000 cells to each treatment.

At the end of the experimental period (28 days), a subset of the specimens from both treatments (n_live_=6-8, n_dead_=10-12) were randomly collected and prepared for MicroCT scanning. Samples of experimentally treated live specimens were bleached in 6% sodium hypochlorite solution (Fisher Scientific NC1796686) for 12 hours to remove organic material from the test. The bleached tests from both live and dead cell treatments were washed twice with Milli-Q water, followed by subsequent washing with 50%, 80%, and 100% ethanol to rinse any remaining debris and dehydrate the tests in preparation for microscopic imaging (**Supplemental Figure S3**). All the live and dead treatment specimens were stored in 100% ethanol at 4°C until microCT scanning.

### Seawater carbonate chemistry

Filtered tank water stored at -20°C was used for carbonate-system analysis. Quality control for pH data was assessed three times per week with Tris standard (Dickson Lab Tris Standard Batch 205) and handheld conductivity probes used for discrete measurements were calibrated once per week. Total alkalinity (TA) was measured using an open-cell titration (48) with certified HCl titrant (∼0.1 mol kg^−1^, ∼0.6 mol kg^−1^ NaCl; Dickson Lab) and TA measurements identified < 1% error when compared against certified reference materials (Dickson Lab CO2 CRM Batch 196). Seawater chemistry was completed following Guide to Best Practices (48). Tri-weekly measurements were used to calculate carbonate system parameters (**Table 1**), using the SEACARB package (49) in R v3.5.1 (R Core Team, 2018).

### Imaging of foraminifera tests with microCT scanning

Foraminifera tests (untreated, dead treated, and live treated) preserved in 100% ethanol were allowed to air dry completely before mounted with Bondic resin (Bondic, CECOMINOD032561) on a flat surface and cured under UV light. Coordinates of mounted tests were identified through a prescan with a Zeiss Xradia Versa 610 X-Ray microscope under the 0.4x objective using the following parameters: 50kV voltage, 4.5W power, 401 projections. Identified tests were individually imaged with the following imaging parameters under the 4x objective: 80kV voltage, 10W power, 2401 projections. Stacked TIFF images were produced based on automatic reconstruction settings during the imaging. The resulting image stacks were imported into the Dragonfly image analysis software (ORS systems Core dll version 2022.2.0.1399, Montreal, CA), which creates a 3D-reconstruction for each foraminifera test. The 8 newest chambers in the 3D-reconstruction of each test were manually isolated through the graphical interface of the Dragonfly software by extracting a region of interest (ROI) containing test areas that are visible from the outside and deleting any undesired regions (e.g., the sutures or air bubbles introduced by the mounting process) (**Figure 1B-C**). Voxels with an intensity lower than 32,000 were filtered out from each chamber, preserving regions that contained the calcium carbonate test, but excluding voxels that imaged the resin or most air bubbles. The extracted ROIs were then used to calculate a thickness mesh using the “generate thickness mesh” function in Dragonfly, where thicknesses throughout the test were calculated by fitting spheres between the outer and inner test surfaces. The thickness mesh of each test chamber was individually exported to a csv file and used for statistical analysis. The number of thickness measurement data points exported ranged between 49,721 and 700,539 per chamber, covering the entire ROI of each chamber.

### Classification of microspheric and megalospheric foraminifera

Diameter of the proloculus and the overall test were determined by fitting a smallest possible sphere over their corresponding outer surfaces using the Dragonfly image analysis software, where the radius of the fitted sphere was reported and used for calculating the diameter of its corresponding proloculus or test. The number of chambers present in each foraminifera was manually counted based on an internal slice projection that included all chambers. During analysis of the 3D-reconstruction of foraminifera tests, a bimodal distribution of proloculus diameters were observed, resulting in two populations: (1) megalospheric, tests with proloculus diameter greater than or equal to 55 μm, (2) microspheric, tests with proloculus diameter less than 55 μm (**Supplemental Figure S1**). Correspondingly, these two populations had distinct distributions of the number of chambers (**Figure 2B**).

### Data analysis

All statistical analysis was performed in R v4.2.3 using the sjstats package version 0.19.0 and the stats package version 4.2.3. Results were visualized in R v4.2.3 using ggplot2 version 3.4.1 and plotly version 4.10.4. The number of chambers per test and the test diameters were compared between microspheric and megalospheric foraminifera using one-way analysis of variance (ANOVA) (**Figure 2B**). Growth of live foraminifera throughout the treatment period was approximated by comparing their number of chambers to the pool of dead treated and untreated specimens (referred to as “not-live”) using two-way ANOVA that accounted for differences in microspheric and megalospheric samples, followed by the Tukey’s honestly significant difference test (TukeyHSD) (**Figure 2C**).

Due to the low abundance of megalospheric specimens in several treatments (**Supplemental Table S3**), all the statistical analyses related to test thicknesses were performed with only the microspheric foraminifera. To normalize the thickness measurements from microCT scanning, test thicknesses were divided by the diameter of each corresponding test. The normalized thickness values were compared using two-way ANOVA that accounted for a treatment factor (e.g., treated versus untreated, different *p*CO_2_ conditions, or live versus dead treatment) and a second factor that accounted for variations of individual foraminifera. To account for the large number of thickness measurement data points from each specimen, all ANOVA analyses that showed statistical significance were followed by the calculation of effect size (η^2^) measures (**Table 2**, **Figure 3**, **Figure 4**). The effect size ranges from 0 to 1 and is representative of the proportion of variance in the model explained by a given factor.

Specifically, test thickness differences between experimentally treated and untreated foraminifera were examined separately with live or dead specimens and across the three *p*CO_2_ treatments (**Table 2**). Variation of test thicknesses across different *p*CO_2_ conditions were compared separately for the live or the dead treatments (**Figure 3A, 3B**), and the variation between live and dead foraminifera were compared separately for the different *p*CO_2_ conditions (**Figure 3C-3E**). Finally, test-thickness variations between live and dead specimens were examined within each of the eight newest chambers (from n to n-7) to assess their differential responses to the different *p*CO_2_ conditions (**Figure 4**).

## Data availability

Data files including water chemistry data and shell thickness measurements are available on figshare at https://figshare.com/s/4464cc33548faf92e211. All scripts used for analysis are available at https://github.com/zhanglab/Foram_OA.

## Acknowledgements

This project was supported by the National Science Foundation Office of Integrative Activities, award #1929078 and an Undergraduate Research Award from the University of Rhode Island (Fall 2022). We thank Dr. Tatiana Rynearson’s laboratory for providing culture of *Skeletonema dohrnii* PA 250716_D1 for the feeding of live foraminifera during acidification treatments.

## Supplemental Figures

**Supplemental Figure S1.**
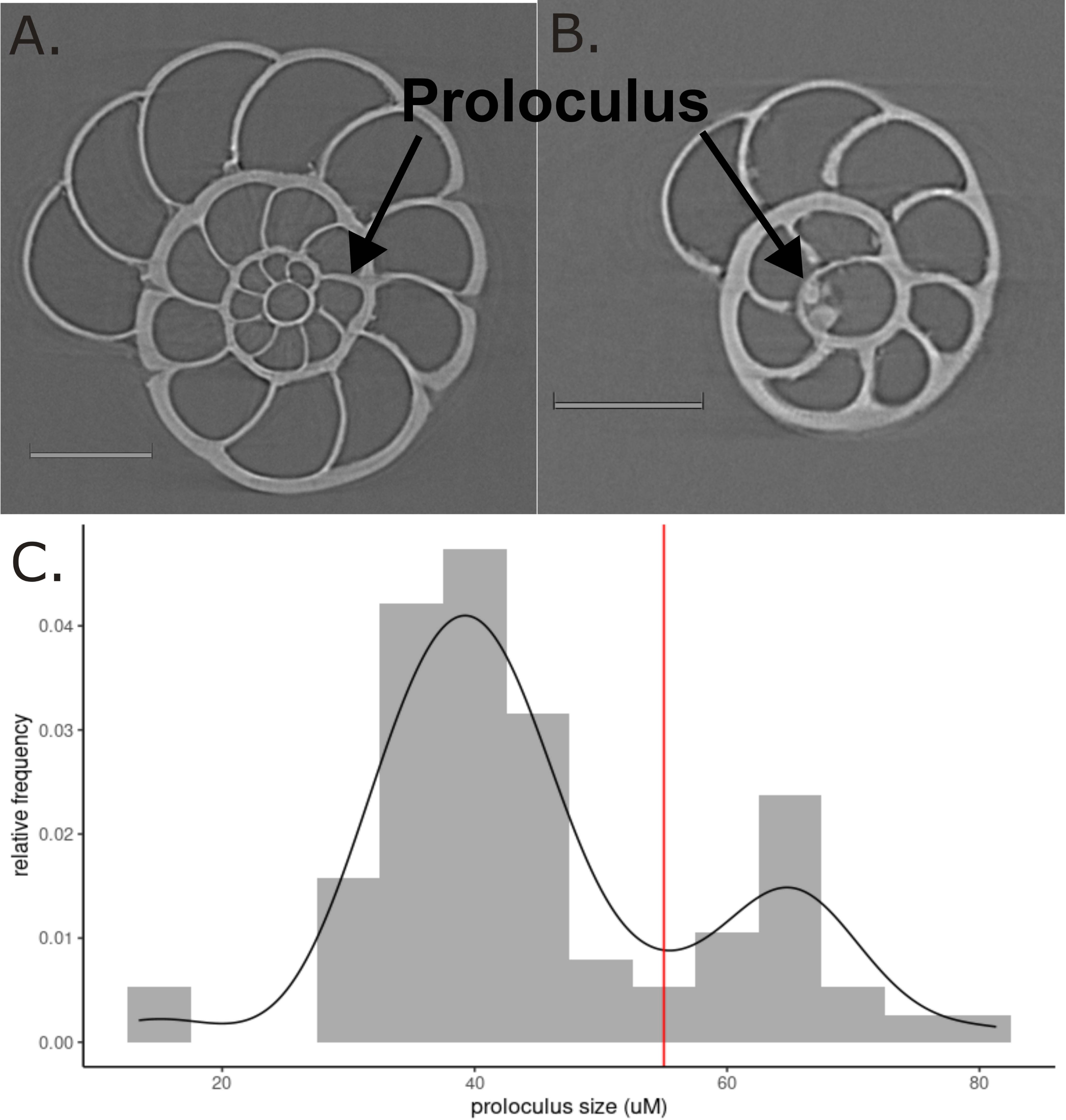
(A) Cross section of a microspheric test. (B) Cross section of a megalospheric test. Each scale bar represents 100 μm. (C) Histogram of proloculus diameters showing a bimodal distribution.

**Supplemental Figure S2.**
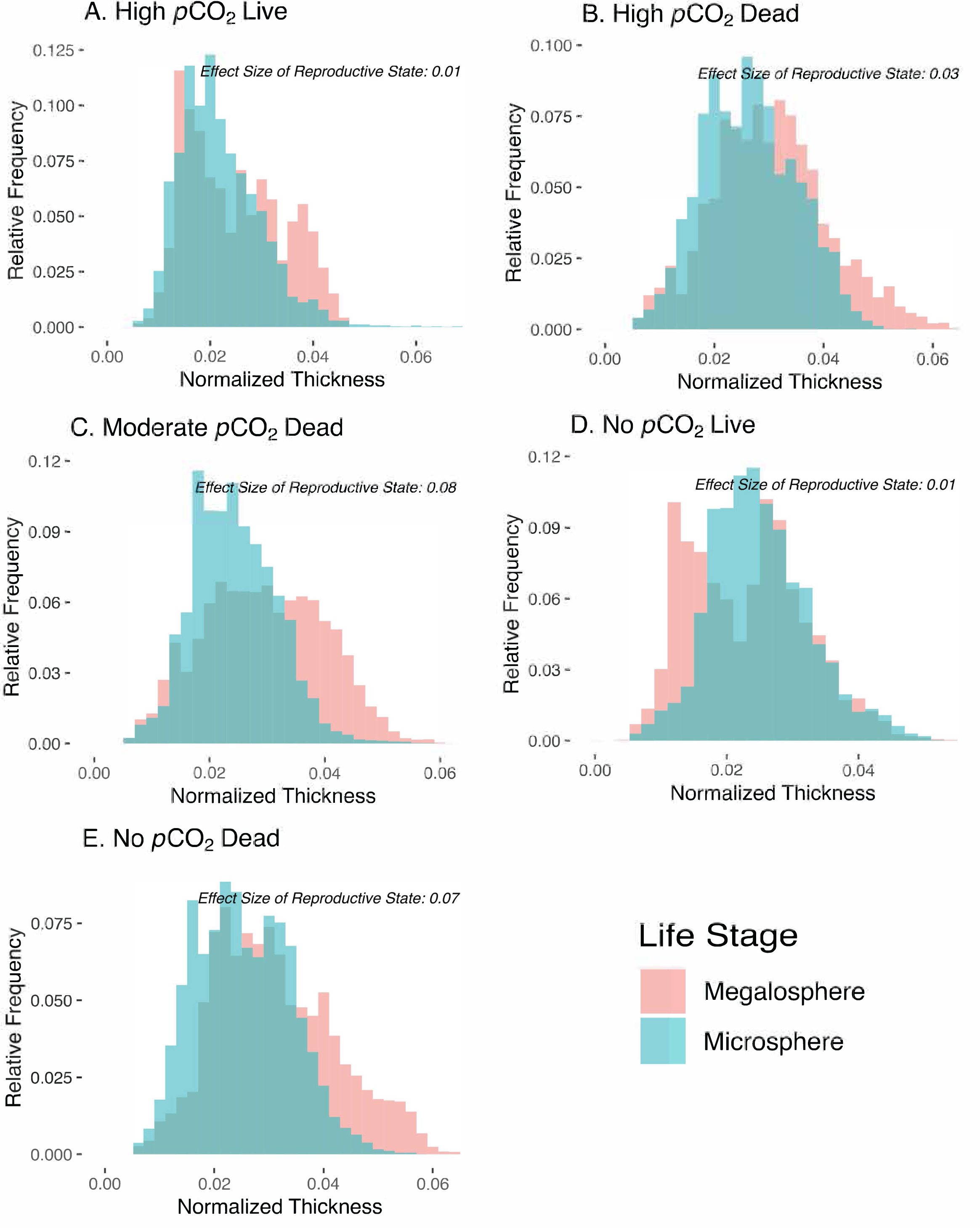
(A-E) Distribution of normalized test thickness between microspheric and megalospheric specimens within each experimental treatment.

**Supplemental Figure S3.**
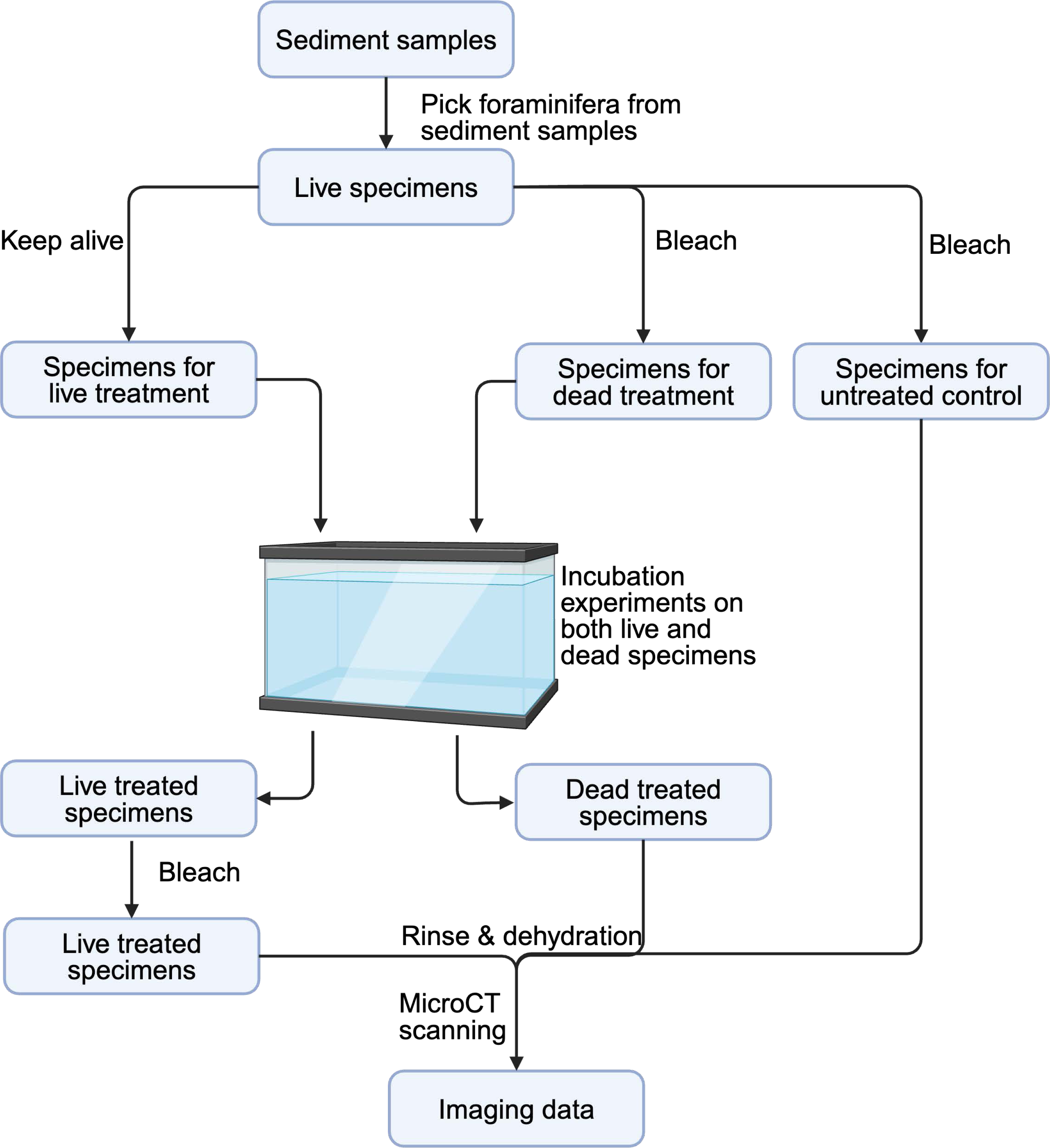
Schematic of experimental workflow as detailed in Materials and Methods.

## Supplemental Tables

**Supplemental Table S1**. Collection of previous ocean and coastal acidification studies of foraminifera, including reference information for the paper, condition of acidification treatments (*i.e.*, pH, *p*CO2), and a summary of key findings.

**Supplemental Table S2**. Tri-weekly water chemistry data from each treatment tank of both the Summer 2023 and Fall 2023 experiments. Parameters shown in the table were calculated using the SEACARB R package V3.5.1 with the exception of pH, temperature, salinity, and total alkalinity (TA), which were experimentally measured.

**Supplemental Table S3**. Number of megalospheric and microspheric specimens collected from the Summer 2023 and Fall 2023 acidification experiments across each treatment. Total indicates the sum of numbers from both replicate experiments.

**Supplemental Table S4**. Measurements of foraminifera test morphology for all specimens analyzed in this study. Data includes the trial number (OA2 or OA3), treatment tank, specimen ID, measurements of test radius/diameter, proloculus radius/diameter, number of chambers, and assignment of foraminifera life stage.

